# Ontogeny and social context regulate the circadian activity patterns of Lake Malawi cichlids

**DOI:** 10.1101/2023.05.27.542564

**Authors:** Evan Lloyd, Aakriti Rastogi, Niah Holtz, Ben Aaronson, R. Craig Albertson, Alex C. Keene

## Abstract

Activity patterns tend to be highly stereotyped and critical for executing many different behaviors including foraging, social interactions and predator avoidance. Differences in the circadian timing of locomotor activity and rest periods can facilitate habitat partitioning and the exploitation of novel niches. As a consequence, closely related species often display highly divergent activity patterns, raising the possibility that a shift from diurnal to nocturnal behavior, or vice versa, can occur rapidly. In Africa’s Lake Malawi alone, there are over 500 species of cichlids, which inhabit diverse environments and exhibit extensive phenotypic variation. We have previously identified a substantial range in activity patterns across adult Lake Malawi cichlid species, from strongly diurnal to strongly nocturnal. In many species, including fishes, ecological pressures differ dramatically across life-history stages, raising the possibility that activity patterns may change over ontogeny. To determine if rest-activity patterns change across life stages we compared the locomotor patterns of six Lake Malawi cichlid species. While total rest and activity did not change between early juvenile and adult stages, rest-activity patterns did, with juveniles displaying distinct activity rhythms that are more robust than adults. One distinct difference between juveniles and adults is the emergence of complex social behavior. To determine whether social context is required for activity rhythms, we next measured locomotor behavior in group housed adult fish. We found that when normal social interactions were allowed, locomotor activity patterns were restored, supporting the notion that social interactions promote circadian regulation of activity in adult fish. These findings reveal a previously unidentified link between developmental stage and social interactions in the circadian timing of cichlid activity.

## Introduction

Animals display remarkable diversity in rest-activity patterns^1,2^. The timing of rest and activity can differ dramatically between closely related species, or even between populations of the same species, raising the possibility that it can be adaptive and subject to selection^3,4^. Indeed, circadian regulation of locomotor activity is strongly associated with many factors critical in determining organismal fitness, including foraging strategy, social behavior, and predator avoidance^5,6^. Further, rest-activity patterns are acutely regulated by environmental factors and life-history traits that include food availability, social interactions, and age^7,8^. Defining how complex environmental interactions regulate activity patterns is therefore critical to understanding behavioral adaptation and evolution.

There are over 35,000 teleost species, adapted to diverse habitats, and representing ∼50% of vertebrate diversity^9^. Many teleosts display robust diurnal locomotor rhythms including the goldfish (*Carassius Auratus*), river-dwelling populations of the Mexican tetra (*Astyanax mexicanus*), and the zebrafish (*Danio rerio*)^10–13^. Examples of nocturnal teleosts have also been identified including the plainfin midshipman, the Senegalese sole, and the doctor fish, *Tinca tinca*^14^. Other species such as cavefish morphs of the Mexican tetra, *A. mexicanus*, and the Somalian cavefish, *Phreatichthys andruzzii*, and cave dwelling populations of *A. mexicanus* have largely lost light-driven circadian regulation of behavior^15–18^. Despite the characterization of species from disparate lineages/populations, few studies have examined how developmental stage, or divergent ecological contexts regulate sleep among closely related species. Understanding the variability of rest-activity patterns over phylogeny, ecology and ontogeny, therefore represents a vital step toward identifying conserved genetic and evolutionary features that may influence regulation of activity throughout vertebrates.

Lake Malawi cichlids exhibit unparalleled diversity in morphology and behavior among vertebrates^19–21^. Observations at night suggest that adult cichlids may be diurnal as an evolved strategy for predator avoidance, at least for species occupying near-shore habitats^22^. Specifically, Lake Malawi is home to endemic non-cichlid predators, including the Cornish jack *Mormyrops anguilloides*, which feeds in packs at night using weak electrical pulses thought to be undetectable by cichlids^23^. We previously analyzed the activity patterns of 11 species of cichlids, from diverse habitats and distinct genetic lineages and found significant variability ranging from highly nocturnal to diurnal, with many species exhibiting no differences in day or night activity^24^. In a single identified nocturnal species, *Tropheops*. sp. “red cheek,’ the pattern held at two life-history stages (i.e., late juvenile vs. mature adult), and under different abiotic conditions (i.e., presence vs. absence of shelter)^24^. However, for many species there was no preference for light or dark activity, raising the possibility that activity patterns are either context dependent or absent in some species^24^.

Here, we focused on the activity patterns of Lake Malawi cichlids that seem to lack activity patterns during adulthood. All Lake Malawi cichlids are maternal mouth brooders, and we found that as newly emerged fry (i.e., early juvenile stage) all six species exhibited robust rest-activity patterns. Thus, we demonstrate ontogenic regulation of activity patterns across diverse cichlids. Furthermore, in a subset of species, we find that activity patterns of adults were largely restored under social housing conditions. Together, these studies demonstrate the complexity and context dependency of the circadian regulation of rest-activity in Lake Malawi cichlids.

## Results

### Total activity is similar between juvenile and adult cichlids

We examined six Lake Malawi cichlids from three distinct lineages (Fig 1A). The *mbuna* lineage is phylogenetically monophyletic, with species generally inhabiting the near-shore rocky habitat. Three *mbuna* species were utilized here: *Labidochromis caeruleus* (Lundo Island), *Melanochromis heterochromis* (Mumbo Island), *Tropheops kumwera* (Kanchedza Island). All three are territorial and sexually dimorphic as adults. *Labidochromis* species are generally omnivorous, consuming both benthic invertebrates, algae, and plankton. *M. heterochromis* has a similarly omnivorous diet. *T. kumwera* feeds primarily on benthic algae^25,26^. *Astatotilapia calliptera* is sister to the *mbuna*, and has been called the “most generalized species in the lake”^26^. It’s generalist designation refers to both diet and habitat, as it one of few Lake Malawi species that inhabits the surrounding river systems. Both *Aulonocara stuartgranti* and *Nimbochromis venustus* are non-*mbuna* species, which is a polyphyletic group of cichlids that generally occupy deeper, open-water, and/or sandy habitats. *A. stuartgranti* feeds on benthic invertebrate that it locates using an enlarged lateral line system, whereas *N. venustus* is an open-water piscivore. All fish were obtained from the aquarium trade. While generation from the wild cannot be verified for these animals, breeding populations are maintained for species, and often to specific locations/populations in the lake, with new individuals introduced to the breeding pool to maintain genetic health.

**Figure 1.**
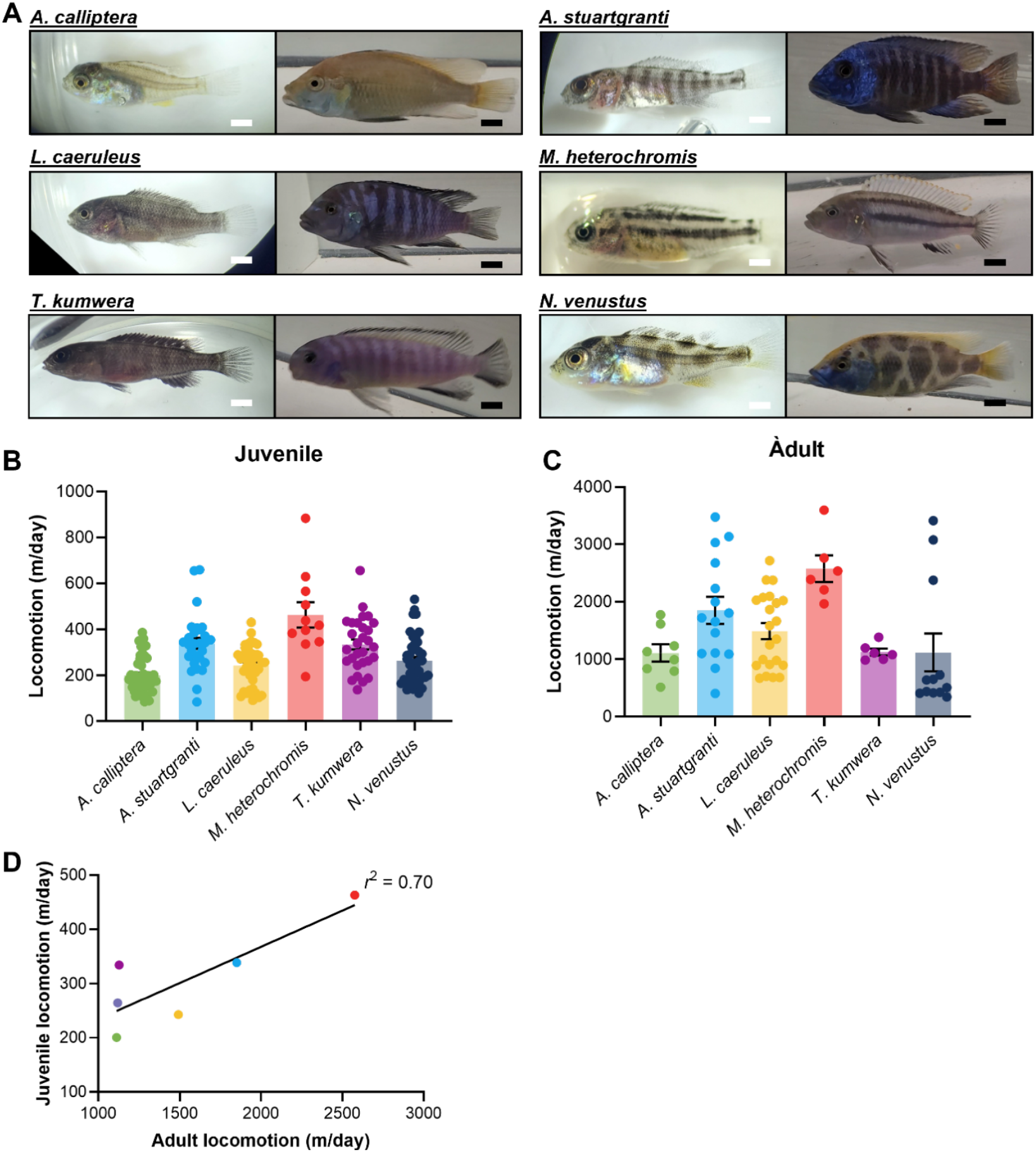
Variation in cichlid activity levels is conserved throughout development. **A**. Images of early juvenile (left) and adult (right) cichlid species used in the present study. Juveniles were tested and photographed between 3-4 weeks post-fertilization, shortly after the depletion of the yolk sac and at the earliest free-swimming stage. Adults were tested and photographed at full maturity, after the development of nuptial colors. White Scale Bar (Juveniles) = 1 mm. Black Scale Bar (Adults) = 1 cm. **B**. Total locomotion of juvenile cichlids over a 24 hour period. There is significant variation in total locomotion in juvenile cichlids (ANOVA: F_5, 190_ = 15.99, *p*<0.0001). **C**. Total locomotion of adult cichlids over a 24 hour period. There is significant variation in total locomotion in adult cichlids (ANOVA: F_5, 67_ = 6.42, *p*<0.0001) **D**. Correlation between 24-hour locomotion in juveniles and adults. There is a significant correlation between juvenile and adult locomotion (*r*^2^ = 0.70, *p* = 0.0388)

To measure activity in cichlids we used systems similar to those established in other fish species including zebrafish and *A. mexicanus*^27–29^. For juveniles, we used 6-well tissue culture plates, whereas adults were filmed in 2.5-gallon glass aquaria. Following 24 hours of acclimation, individually housed fish were filmed for a 24 hour period under standard 14 hr light: 10 hr dark conditions. We noted significant variation in total activity with and between species of juvenile and adults (Fig. 1B, C). For example, *A. calliptera* was the least active in both juveniles and adults, while *M. heterochromis* had the highest activity levels at both stages, suggesting total activity is conserved across developmental stages (Fig 1B,C). To directly test this notion, we examined the relationship between total activity in juveniles and adults. A regression revealed a strong positive relationship (r^2^=0.70, *p*=.038) across species between juvenile adult locomotor activity (Fig 1D). Together, these data reveal significant inter-species variation that remains constant across the life cycle.

### Juvenile cichlids exhibit robust activity patterns compared to adults

We next sought to understand whether rest-activity patterns changed between life-history stages. In juveniles, we identified robust activity patterns across all six species tested (Fig 2A,C). In total there were four diurnal species (*A. calliptera, L. caeruleus, M. heterochromis*, and *T. kumwera*) and two nocturnal species (*A. stuartgranti* and *N. venustus*). Conversely, there were no significant differences between day and nighttime activity across all six species when tested during adulthood (Fig 2B,D). These results were confirmed when we computed a diurnality index, which compares daytime to nighttime activity in individual fish (Fig 2E)^24^. There was significantly more time-of-day activity preference in five of the six species tested (Fig 2E). Thus, activity patterns appear to be more robust in early juvenile cichlids compared to adult fish under individually housed conditions.

**Figure 2.**
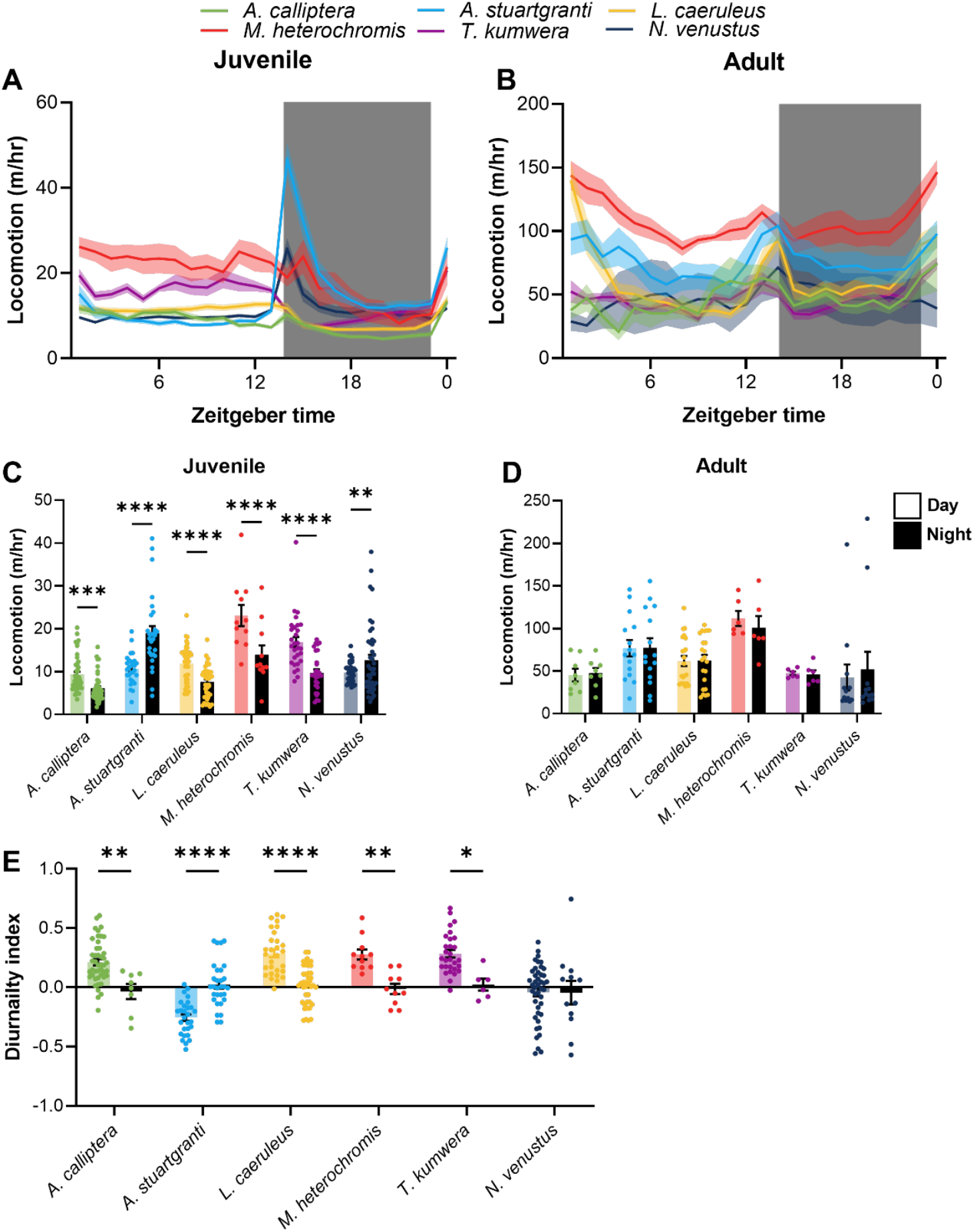
Ontogeny of behavioral rhythms in locomotion. **A**. Locomotion profiles of juvenile cichlids over 24 hours. Shaded area indicates dark period. **B**. Locomotion profiles of adult cichlids over 24 hours. Shaded area indicates dark period. **C**. Average hourly locomotion of juvenile cichlids during the day and night. Juvenile cichlids exhibit diurnal (*A. calliptera, L. caeruleus, M. heterochromis, T. kumwera*) or nocturnal (*A. stuartgranti, N. venustus)* activity patterns (two-way ANOVA: F_5, 190_ =36.54, *p*<0.0001). **D**. Average hourly locomotion of juvenile cichlids during the day and night. Adult cichlids lack behavioral rhythms when tested in isolation. **E**. Strength of behavioral rhythms in juvenile and adult cichlids. 1 indicates total diurnality, -1 indicates total nocturnality. Behavioral rhythms are present in juveniles but not adults (two-way ANOVA: F_5,280_ = 16.20, *p*<0.0001).

Across taxa, the timing of behavioral quiescence or rest are modulated by the circadian clock and linked to daily activity patterns^3,4^. To examine whether the timing of rest differs across life stages, we therefore compared rest periods of one minute or longer, a timeframe of inactivity that is used to define sleep in related species^30^. There were robust differences in total rest across juvenile and adult species (Fig 3A-D). Similar to the analysis of activity, there was consistency in the duration of rest between juveniles and adults. For example, *M. heterochromis* had the lowest levels of rest in juveniles and adults, whereas *A. calliptera* and *N. venustus* had the highest level at both developmental stages. Indeed, a regression analysis revealed a strong relationship (r2=0.82, p=0.012) between juvenile and adult rest duration (Fig 3E). Therefore, the duration of total rest is maintained throughout development across multiple species of cichlids.

**Figure 3.**
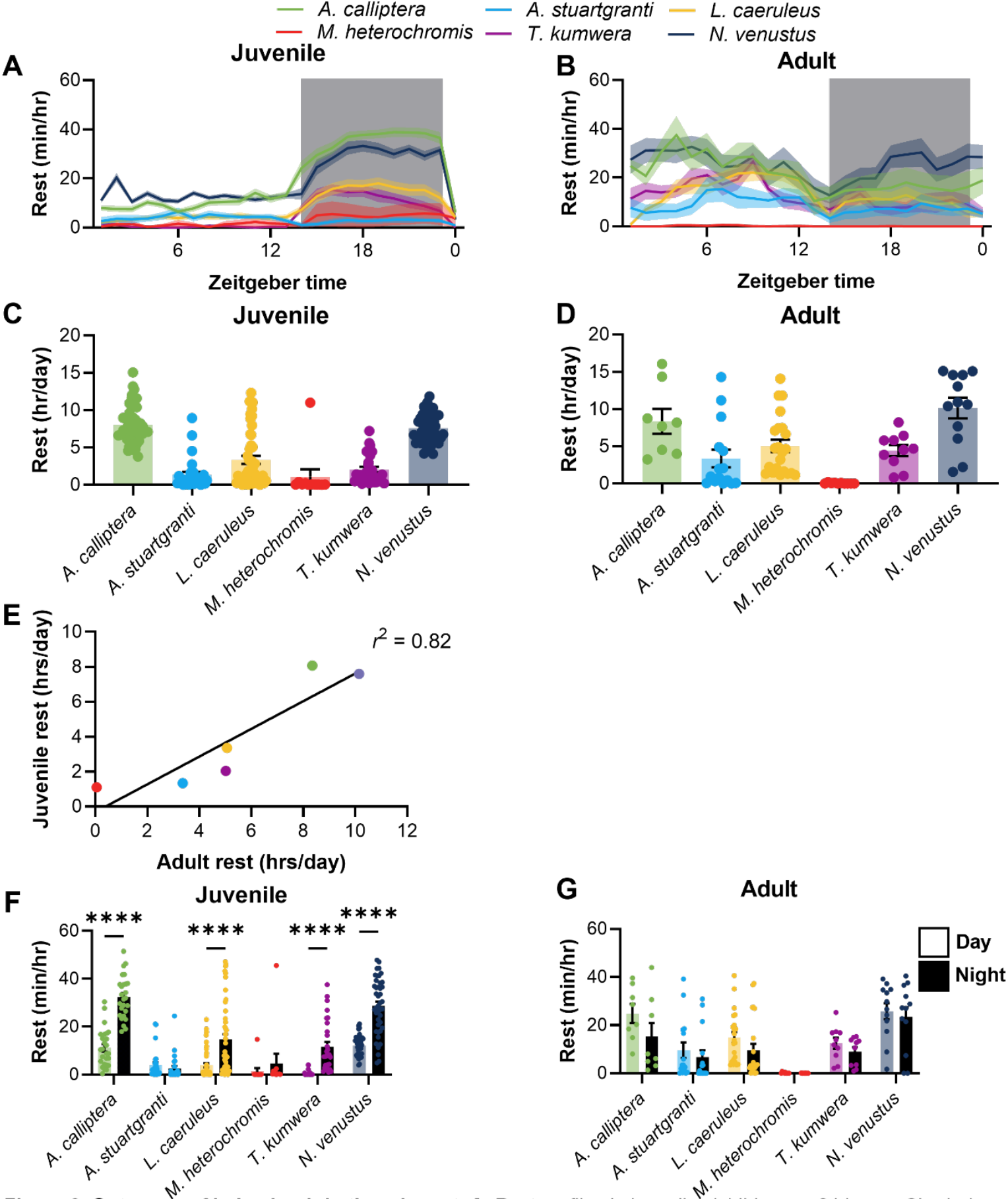
Ontogeny of behavioral rhythms in rest. **A**. Rest profiles in juvenile cichlids over 24 hours. Shaded area indicates dark period. **B**. Rest profiles in adult cichlids over 24 hours. Shaded area indicates dark period. **C**. Total rest amounts over 24 hours in juvenile cichlids. There is significant variation in total rest amount in juvenile cichlids. (ANOVA: F_5, 190_ = 42.4, *p*<0.0001). **D**. Total rest amounts over 24 hours in adult cichlids. There is significant variation in total rest amount in adult cichlids (ANOVA: F_5, 67_ = 8.792, *p*<0.0001). **E**. Correlation between juvenile and adult rest amounts. There is significant correlation between juvenile and adult rest (*r*^2^ = 0.82, *p*<0.0128). **F**. Average hourly rest amount of juvenile cichlids during the day and night. Juvenile cichlids exhibit diurnal (*A. calliptera, L. caeruleus, T. kumwera, N. venustus*) or nocturnal (*A. stuartgranti)* rest patterns (two-way ANOVA: F_5, 358_ =10.95, *p*<0.0001). **G**. Average hourly rest amount of adult cichlids lack behavioral rhythms in rest amount.

We next compared the amount of rest over the light and dark periods. Juvenile *A. calliptera, L. caeruleus*, and *T. kumwera* displayed robust patterns in rest regulation with increased nighttime rest, while the fourth diurnal species *M. heterochromis* trended in the same direction (Fig 3F), suggesting diurnality is associated with periods of nighttime rest in juvenile cichlids. Conversely, there were no differences in the timing of rest for *A. stuartgranti*, suggesting that nocturnality in this species is driven by swimming velocity during wakefulness, rather than the timing of rest (Fig 3F). Finally, in *N. venustus*, which is nocturnal at the junvenile stage, there is robust consolidation of rest during the night (Fig 3F). For all species tested there was little difference between daytime and nighttime rest in adults (Fig 3G). To understand this trait, we calculated a crepuscularity index, which measures the ratio of activity immediately after the light transitions relative to the rest of the day. We saw notable variation across species in this measurement, with *N. venustus* and *A. stuartgranti* displaying the highest degree of crepuscularity, increasing activity levels by up to two fold in the hours following light transitions; other species showed little to no crepuscular tendencies (Fig S1A). Similar to day-night behavioral rhythms, crepuscular behavior was largely lost in adulthood (Fig S1B).

There were no day-night differences in rest across all six populations of adult cichlids, again confirming that the timing of rest-activity is more robust in juveniles than adult fish (Fig 3G). Together, these findings support the notion that rest-activity bouts are more readily consolidated in juvenile fish and reveal separate mechanisms for the emergence of diurnal, nocturnal, and crepuscular activity patterns.

### Social housing can restore diurnal activity in adults

Many fish species, including cichlids, are highly social, with animals displaying behavioral differences under solitary and group housing ^31–35^. Except for a small number of studies, nearly all analysis of rest-activity patterns in fishes have used individually housed animals, including cichlids. Social interactions are weaker in juvenile fish, compared to adults, raising the possibility that the lack of rest-activity patterns in adults is an artifact of solitary housing conditions. To examine the effects of social housing on rest-activity patterns we measure activity patterns in group housed cichlids comprised of two adult males and four adult females (Fig 4A). All fish were transferred into testing tanks and given 24 hours to acclimate and establish stable social interactions. We tested four different cichlid species under group-housed conditions for 24 hours. While we were unable to track individual fish reliably over the 24 hour period, we used average activity of the group. Three of these species tested (*A. calliptera, L. caeruleus*, and *T. kumwera*) trended towards recovery of diurnal behavior (Fig 4C). There was little difference in the species *A. stuartgranti* or *N. venustus*, which are nocturnal as juveniles. To examine the broader trends of sleep in group housed fish we combined analysis of nocturnal and diurnal species. Social housing induced robust diurnality in species that are diurnal as juveniles, while there was no differences between day and night activity in the two nocturnal species (Fig 4D). These findings suggest diurnal juveniles species likely maintain activity patterns in adulthood, but the fish only display these behaviors under group-housed conditions.

**Figure 4.**
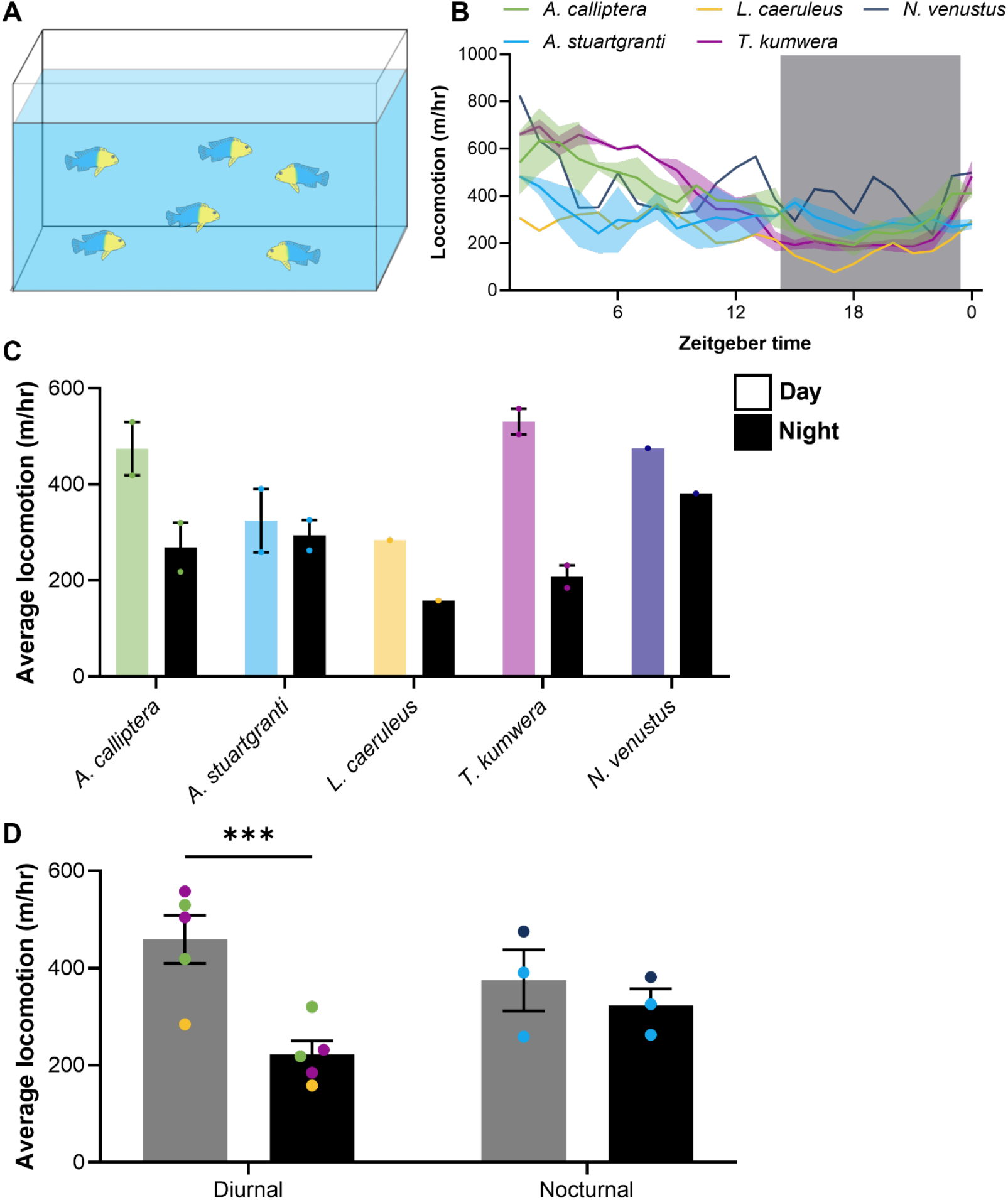
Social context restores rhythms in diurnal cichlids. **A**. Schematic of test setup. Adult cichlids are transferred to a 10-gallon testing aquarium, acclimated overnight, and then recorded for 24 hours. **B**. Average locomotion of cichlid groups over 24 hours. Shaded area indicates dark period. **C**. Average hourly locomotion of group housed cichlids, during the day and night. **C**. Average hourly locomotion of group housed cichlids during the day and night, grouped into diurnal and nocturnal species. Group housing restores rhythmic behavior in diurnal, but not nocturnal species (two-way ANOVA: F_1,6_ = 11.23 *p* = 0.0154). Colors of data points indicate species.

To examine the behavior of individual fish in more detail, we manually annotated behavior for a single species over the full 24 hour testing period. We chose to examine *L. caeruleus* because of their robust diurnal behavior as group-housed adults and juveniles. Fish behavior was manually analyzed in Behavioral Observation Research Interactive Software (BORIS) for active swimming over the 24 hour cycle^36^. We developed an ethogram across all six fish with rest-activity timing (Fig 5A). We observed robust diurnal activity across all male and female individuals (Fig 5B,C). Further, the duration of rest bouts was greater during the night period (Fig 5D,E). These findings confirm that diurnality is not sex-specific, and generalizable across the dominance structure.

**Figure 5.**
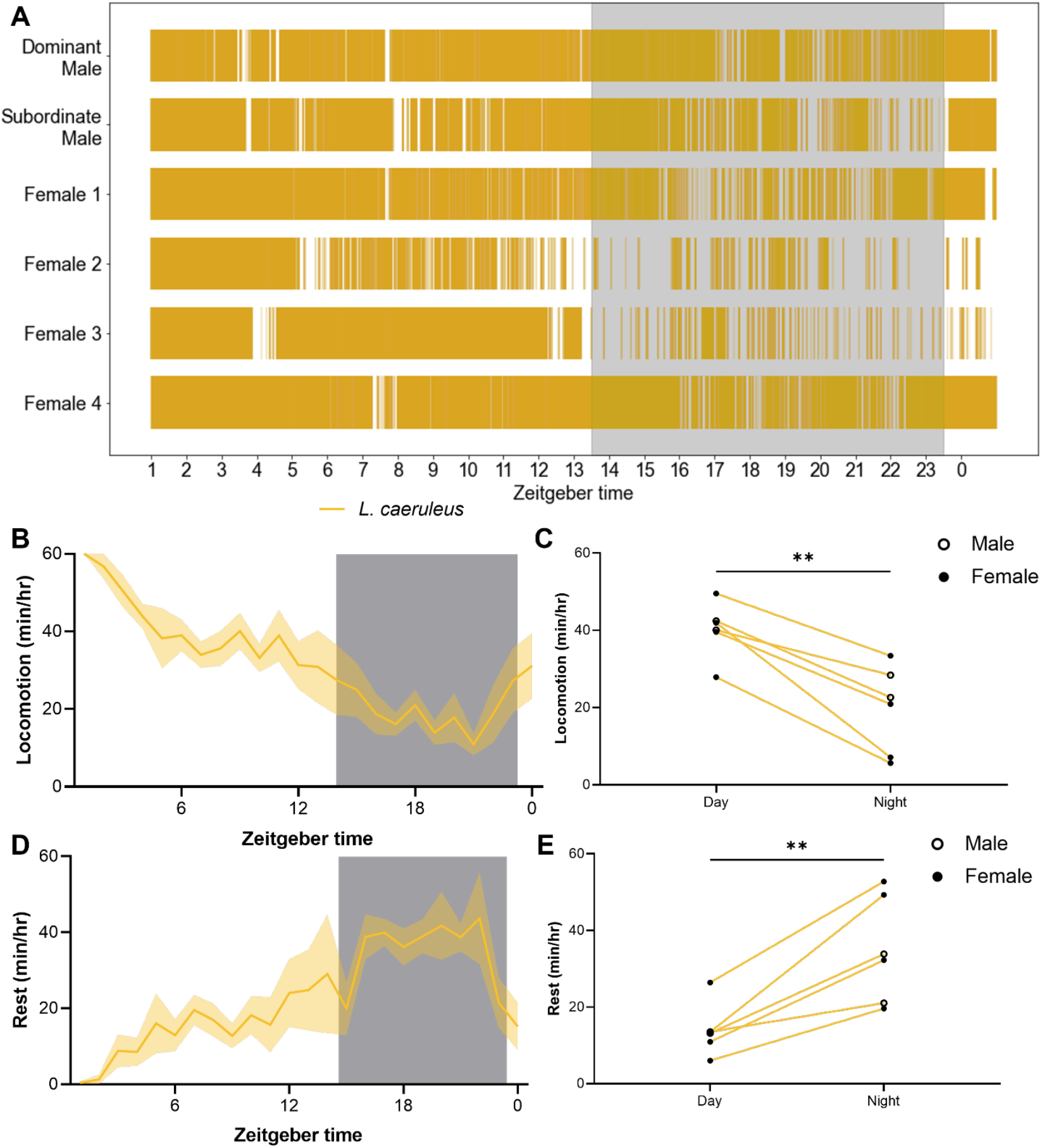
Individual variation in behavioral rhythms in a social context. **A**. Ethogram of active swimming behavior of *L. caeruleus* over 24 hours, in a social context. Colored regions indicate periods of active swimming. Shaded area indicates dark period. **B**. Profile of locomotion in group housed *L. caeruleus*, quantified as minutes of active swimming per hour. Shaded area indicates dark period. **C**. Average hourly locomotor activity of group housed *L. caeruleus* during the day and night. Group housed *L. caeruleus* are significantly more active during the day (Paired t-test: t_5_ = 6.413, *p =* 0.0014). **D**. Rest profile of group housed *L. caeruleus* over 24 hours. Shaded area indicates dark period. **E**. Average hourly rest of group housed *L. caeruleus* during the day and night. Group housed *L. caeruleus* consolidate rest during the dark period (Paired t-test: t_5_ = 5.198, *p* = 0.0035).

## Discussion

### The importance of context in behavioral studies

The genetic, morphological, and behavioral diversity of African cichlids provides an exceptional model for understanding trait evolution^19^. In spite of myriad comparative studies in this system, very few have focused on rest-activity patterns. Here, we characterize the behavior of six species of Lake Malawi cichlids at multiple developmental stages, highlighting a role for ontogeny and social context in the regulation of activity. Previously, we characterized rest-activity patterns in 11 species of Lake Malawi cichlids at the late juvenile, or sub-adult stage, identifying 7 species without activity preferences, as well as two diurnal species and one nocturnal species^24^. The lack of activity preferences across the majority of species was surprising given the ubiquity of circadian rhythms and the robust diurnal or nocturnal behaviors across many species of animals that have been studied to date. Here, we examined species that also seemed to lack robust activity patterns as adults, but showed that these behaviors are present when animals were examined at an early juvenile stage or within a social context. Of note, *T. kumwera* showed diurnal activity patterns in our previous study, when tested at the sub-adult stage. Here, we found that this behavioral rhythm is present from the earliest developmental timepoints, but is lost by the onset of sexual maturity. However, placing the adult fish in a social context restored diurnal rhythmicity, underscoring the importance of a social context for maintenance of behavioral rhythms in this species. These findings highlight the complexity of animal behaviors, and suggest that many of the species previously examined may possess innate activity rhythms when examined in a different context.

The role of circadian and sleep regulators across development remains poorly understood. For example, in humans sleep varies significantly across the life cycle but little is known about the factors that regulate pediatric sleep^37^. In fruit flies, genes regulating sleep and activity in larvae differ from those regulating sleep in adult flies suggesting that the genetic basis of the consolidation of activity patterns is specific to developmental stages^38,39^. Here we find that total locomotor activity and duration of rest appear to be robust to life-history stage, whereas the timing of rest-activity patterns is dependent upon ontogeny, raising the possibility that there are genetic and contextual differences regulating the timing, but not the amount of activity. The cichlid species examined here reach sexual maturity at ∼6 months. This stage is associated by an increase in sex-specific hormones that is associated with aggression, territoriality, mating behaviors, and dominance hierarchies^40^. The gain of these behaviors seems to be associated with the loss of activity pattern robustness, suggesting that natural social conditions are required for the expression of diurnal or nocturnal rhythms. Whether the lack of rhythms in individually housed adults is due to the lack of a necessary environmental cue, or represents a secondary stress response remains unclear. Taken together, our data suggests that understanding the rest-activity patterns in adults will require further testing in diverse ethological contexts.

It is notable that total rest-activity appears to be robust to life-history stage, which suggests that it is more hard-wired compared to rest-activity patterns. If true this would suggest greater potential to modulate activity rhythms as populations face new environmental and/or social contexts. Our findings suggest the evolution of highly disparate locomotor patterns, diurnality and nocturnality, are influenced by environmental/social context. These findings raise the possibility that the evolution of complex social behaviors co-evolve with rest-activity patterns to contribute to the regulation of circadian behavior.

### Biological mechanisms of circadian regulation

The hypothalamus is the primary regulator of circadian behavior and outputs from the suprachiasmatic nucleus are thought to convey diurnal or nocturnal behavior^41,42^. In zebrafish, the SCN contains core-transcriptional clock genes common in mammals suggesting a conserved function in the regulation of locomotor patterns, although the presence of many light-responsive tissues in teleosts suggests clock regulation may be more complicated than in mammals^41^. In *Drosophila* through mammals social interactions contribute to the regulation of sleep and circadian behavior. In mammals, social interactions impact clock entrainment. For example, social defeat decreases the amplitude of the core-clock gene Per2 in the periphery, as well as sleep homeostasis^43,44^. There is reduced complexity of social interactions in young juvenile fish that displayed robust rhythms^35,45^. However, we find that activity patterns are absent in individually housed adult fish. These findings raise the possibility that unlike mammals, stress that impairs circadian function comes from solitary housing, rather than social stress. Indeed, in *Drosophila* solitary housing is associated with sleep dysregulation and disruption of circadian gene expression^46,47^. Understanding both the SCN outputs, and how they are modulated by social context provides a mechanism for understanding how context-dependent regulation of activity patterns evolved. Our findings raise the possibility that these outputs are modulated by social circuits, or other context-specific regulators of behavior.

### Adaptive significance

Development of diversity in behavioral rhythms in cichlids may be a type of habitat partitioning, with cichlid species altering the timing of their behaviors to take advantage of reduced competition for resources such as food and territory^24^. The presence of larger nocturnal predators in Lake Malawi, such as the Cornish Jack *Mormyrops anguilloides*, presents a constraint to this adaptation, requiring nocturnal cichlids to develop strategies for avoiding predation at night^23^. It has been previously reported that *A. stuartgranti* possesses widened lateral line canals which enable them to detect prey in the dark, but this same adaptation may also enable detection and evasion of predators during the dark period^48^. It is interesting that both nocturnal species are also crepuscular, and both are predatory. There is evidence that predator-prey interactions are highest during the twilight hours, and crepuscularity may be an adaptation in predators to facilitate access to both day-active and night-active prey^49,51^. Also notable is that neither species recovered rhythmicity when placed in a group context. This may reflect differences in the level of social interaction in these fish in the wild, such that it is not group context, but rather a yet unidentified factor that is required for adult activity rhythms. It is also possible that rest-activity rhythms are truly lost in these species over ontogeny. In short, there is still much to learn about the regulation of activity patterns in this system, and testing hypotheses about the adaptive significance of these behaviors will require the application of phylogenetic methods on a much larger sampling of taxa.

### Challenges and Future Directions

It has been found that changes in not just light quantity, but also composition (color), occur during twilight, and that the circadian clock machinery is responsive to this change^50,52^.. Because our study used constant levels of white light throughout the day, we were unable to evaluate the effects of changes in light quality on behavioral rhythms; future studies may address this question by using simulated dusk/dawn transitions.

The duration of circadian experiments provide a significant challenge for analyzing data across multiple species. Here, we used multiple approaches to quantify locomotor activity in group-housed fish. First, we used Ethovision to track total activity. While this system is capable of tracking individual fish in a shared arena, the tracks regularly cross-over preventing measurements of activity in individuals over a 24 hour period. To identify the activity of individuals we visually quantified behavior using BORIS. While this approach allows for reliable behavioral measurements in individual fish it lacks precise quantification. Given the time-consuming nature of the analysis, we only analyzed a single experiment of six fish. Recent development in automated tracking including DeepLabCut and IDTracker may provide applications for long-term automated tracking^53,54^. These systems have been effectively applied to measure social behaviors and activity in populations of animals ^53,55^. Further, we have tracked large groups of *A. mexicanus* to examine social interactions^32^. While these applications have yet to be applied to long-term analysis of rest-activity patterns the rapid improvements in processing speed and accuracy are likely to allow for analysis of sleep and circadian regulation of activity under social contexts.

It is widely recognized that rest-activity patterns are regulated by many life-history and environmental traits^56,57^. The growing use of animal models in rest-activity regulation has provided unprecedented insight into the genetic, neural and evolutionary processes that govern these behaviors. However, the vast majority of studies still test animals under individually housed conditions because it provides a simpler method of data acquisition and the removal of variables that may impact behavior. The results of this study suggest that developmental stage and environmental conditions can have profound effects on behavioral regulation, and highlight the need for a more thorough investigation of social influences and other factors that may play a role in regulation of behavior.

## Materials and Methods

### Fish husbandry^37,38^

Cichlids used for experiments were reared following standard protocols approved by the Texas A&M University Institutional Animal Care and Use Committee. Cichlids were housed in the Keene fish facilities at Texas A&M University at a water temperature of 28.5°C, on a 14 h:10h light:dark cycle. Adult cichlids were fed TetraMin tropical flakes (TetraMin) twice a day. Juvenile cichlids were fed live *Artemia* brine shrimp twice daily.

### Fish breeding

Breeding was facilitated by the inclusion of clay pots in the cichlid’s home tanks, which provided arenas for mating behaviors. Breeding occurred spontaneously, and females were periodically visually inspected for an enlarged buccal cavity, evidence of fertilized eggs. For experiments testing the behavior of naturally emerged juveniles, mouthbrooding females were isolated and their tank was checked daily for the emergence of juveniles. Upon emergence, juveniles were transferred to the experimental setup immediately. For all other experiments, mouthbrooding females were allowed to carry their embryos until the hatching stage (approximately 3-5 days). A waiting period prior to extraction increased survival rates of the offspring in our hands.

To extract fertilized eggs, the mouthbrooding female was transferred to a holding tank and briefly restrained by hand, while the mouth was gently opened with the pad of the thumb. The buccal cavity was gently stroked to facilitate egg removal. Following extraction, the female was returned to her home tank, and the eggs were transferred to a 1000mL capacity Erlenmeyer flask (VWR, 10545-842) with an air stone bubbler to maintain oxygen levels. Eggs were left to develop until free-swimming, and the yolk mostly depleted (approximately 3-4 weeks), at which point they were transferred to the experimental setup immediately.

### Behavior measurements

For experiments testing locomotor activity in juvenile fish, juveniles were transferred to individual wells of 6-well culture plates (Falcon, 353046), and placed on a light box constructed of white 1/8” high-density polyethylene plastic (TAP Plastics), which allowed for even diffusion of light to facilitate automated tracking. Light boxes were lit from below with infrared light, using 850 nm LED strips (Environmental Lights). Fish were acclimated to the testing chambers overnight; behavioral recording began the following day at ZT1, and ran for 24 hours. Juveniles were filmed from above, at 15 frames per second with a USB camera (LifeCam Studio 1080p, Microsoft), modified to remove the IR-blocking filter, and with an IR-pass filter (Edmund optics, 43-948) added to ensure consistent lighting during the light and dark periods.

For experiments testing locomotor activity in isolated adult fish, adults were transferred to 2.5 gallon tanks (Carolina Biological Supply, 671226), fitted on the floor and walls with custom-cut white corrugated plastic (3mm thick), allowing for even diffusion of light to facilitate automated tracking. Tanks were lit from behind with infrared light. Fish were acclimated to the testing chambers overnight; behavioral recording began the following day at ZT1, and ran for 24 hours. Adults were filmed from the side, with the same recording equipment described above.

For experiments testing locomotor activity in group-housed adult fish, adults were transferred to 10 gallon tanks (Carolina Biological Supply, 671230), fitted on the floor and walls with custom-cut white corrugated plastic, and lit from behind with infrared light. Fish were acclimated to the testing chambers overnight; behavioral recording began the following day at ZT1, and ran for 24 hours. Each group consisted of 2 male and 3-4 female conspecifics.

### Automated behavioral tracking

Acquired videos were processed in Ethovision XT 15 (Noldus), and positional data over the 24 hour period was extracted and analyzed using a custom-made Python script (v 3.11.3) to calculate locomotor activity on an hourly basis. Analysis of rest patterns was carried out as previously described^24^, with a threshold of 4 cm/s for adults, and 12 cm/s for juveniles. Bouts of inactivity greater than 60 seconds were considered rest.

For group housed experiments, locomotor activity was calculated using Ethovision’s Social Interaction Module (Noldus). Due to limitations of the software, the identities of individual fish could not be maintained over the 24 hour period, so the average locomotion within each group was used for subsequent analysis.

### Manual behavior scoring

To assess individual behavioral rhythms within group housed cichlids, a single 24-hour video was manually scored in BORIS, an interactive behavior logging software^36^. Active swimming behavior was logged as a state event (on/off), according to the following criteria: “Focal fish can be seen actively swimming with appendicular movements even though the pace may change throughout the duration of one bout of movement”. A full 24 hours was logged for each fish (N=2 males, N=4 females) in a single video. The identity of the dominant male was determined by his larger size and nuptial coloration pattern. The females were also identified by size, with the largest female designated “Female 1” and the smallest “Female 4”. A single scorer performed all manual activity logging, to avoid inter-individual differences in evaluation of criteria. Data for each fish was exported in the “Aggregated Events” format, and then analyzed using a custom-made Python script (v3.11.3) to extract hourly measurements of locomotion. Rest bouts were calculated from the aggregated movement data, with any period of inactivity greater than 60 seconds considered a rest bout.

### Statistical Analysis

To identify differences between multiple conditions, such as activity in the light versus dark, or between juvenile and adult behavior, a two-way ANOVA was carried out, and followed by Šidák’s multiple comparisons post hoc test. All statistical testing was carried out using InStat software (GraphPad Prism 9.5).

Diurnality was calculated as described previously, as an “activity change ratio”, with 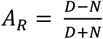 where D and N are average hourly activity during the day and night. Crepuscularity was calculated as 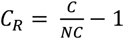, where C is equal to the average activity in the hours following light transitions (i.e. dawn and dusk), and NC is equal to the average activity across the rest of the day. In this formulation, 0 represents no change in activity following a light transition, and 1 represents a 100% increase in activity levels^24^.

## Supplemental Figures

**Supplemental Figure 1.**
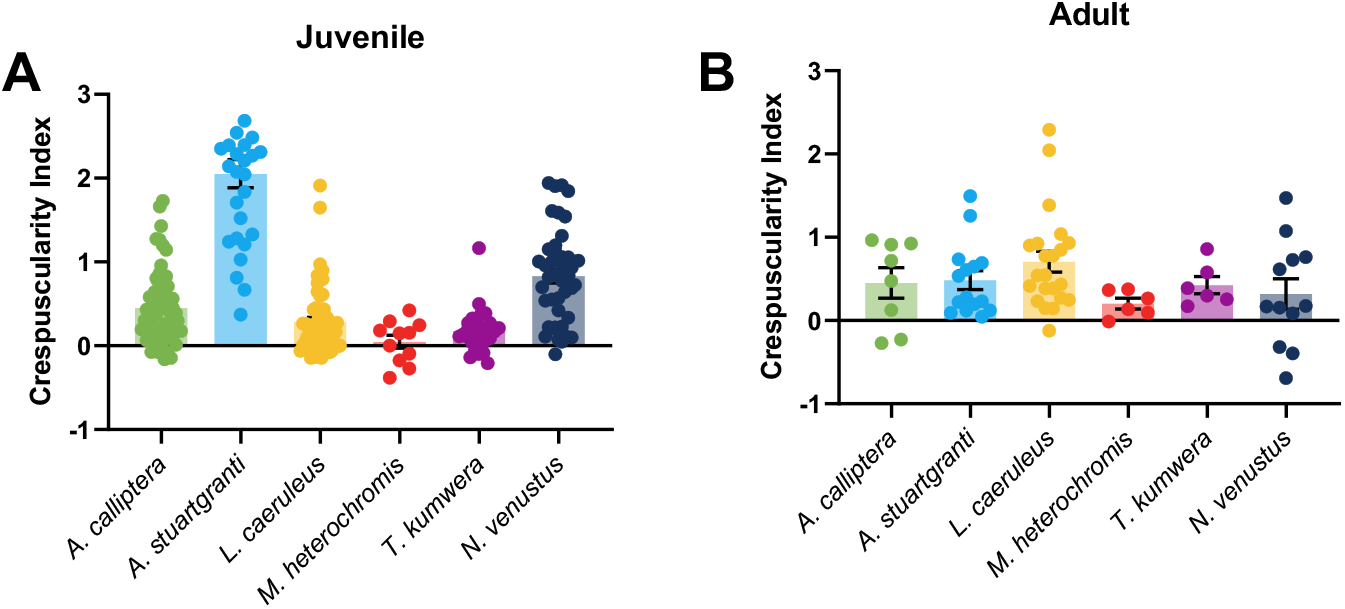
Ontogeny of crepuscular behavior. **A**. Crepuscularity index of juvenile cichlids. There is significant variation in degree of crepuscular behavior among juvenile cichlids (ANOVA: F_5, 190_ = 55.04, *p<*0.0001). **B**. Crepuscularity index of adult cichlids. There is no significant variation in degree of crepuscular behavior in adult cichlids.

